# A Novel Unorthodox Dimeric Primary Enoyl-CoA Reductase Structure

**DOI:** 10.1101/2025.10.17.683039

**Authors:** Cahine Kulakman, Irimpan I. Mathews, Yasuo Yoshikuni, Soichi Wakatsuki, Hasan DeMirci

## Abstract

Enoyl-CoA carboxylases/reductases (ECRs) are enzymes with the fastest carbon dioxide (CO_2_) fixation capabilities, yet the precise mechanisms behind their assembly and catalytic activity are structurally not yet fully understood. Here, we employed cryo X-ray crystallography to reveal the dimeric structural organization of a novel ECR, isolated from mesophilic *Mesorhizobium metallidurans* (*M. metallidurans*). We examined the interactions *in silico* and compared oligomerization of our dimeric ECR from *M. metallidurans* (ECR^Mm_Dim^) with tetrameric ECR from *Burkholderia ambifaria* (ECR^Ba_Tet^)by using size exclusion chromatography in solution. Our *in silico* analysis revealed that specific residues in the *M. metallidurans* ECR that preclude tetramer formation, which could affect the enzyme’s catalytic activity. Additionally, we compared primary ECR sequences and structural variations between *K. setae* and *M. metallidurans* to explore their evolutionary relationships, along with their functional diversity. Our study presents the first example of a dimeric ECR structure which may provide new insights into how dimerization versus tetramerization may have an impact on catalytic function. By detailing how different oligomeric states influence enzyme activity and exploring active site conformational changes, we may offer a further understanding of ECR assembly. This work paves the way for future research into the precise molecular mechanisms that drive ECRs exceptional overall catalytic activity, efficiency and efficacy.

## 1. Introduction

Carbon dioxide (CO_2_) is the primary greenhouse gas contributing to climate change (<u>Cassia et al. 2018</u>, <u>Montzka et al. 2011</u>). Finding efficient ways to convert atmospheric CO_2_ into valuable chemical compounds is of utmost importance in mitigating its environmental impact along with advancing sustainable technologies. Biological CO_2_ fixation, achieved by a variety of enzymes like carboxylases (<u>Aleku et al. 2021</u>), plays a crucial role in the global carbon cycle. Therefore, the focus has shifted from studying natural CO_2_ fixation processes to converting CO_2_ into organic compounds that can be used in diverse applications, ranging from agriculture to renewable energy (<u>Schwander et al., 2016</u>).

A catalytically efficient candidate in this criterion, enoyl-CoA carboxylases/reductases (ECRs) (<u>Erb et al.,</u> <u>2009</u>), has been steadily gaining attention due to its critical role in carbon fixation. ECRs belong to a diverse family of enzymes involved in various metabolic pathways, including the synthesis of fatty acids and dicarboxylic acids through the reduction of α,β-unsaturated enoyl-CoAs, utilizing Nicotinamide Adenine Dinucleotide Phosphate (NADPH) as a cofactor (<u>Erb et al., 2007</u>; <u>Schada von Borzyskowski et</u> <u>al., 2013</u>). Among CO_2_-fixing enzymes, ECRs are notable for their ability to catalyze carbon-carbon bond formation with high speed, specificity, and efficiency, which surpasses many other enzymes in their class (<u>Schwander et al., 2016</u>). Their speed and efficiency highlight the potential of ECRs in the development of innovative biocatalysts for industrial CO_2_ capture and conversion.

Despite their promising capabilities, significant gaps remain in our understanding of the molecular mechanisms underlying ECR function, despite progress in recent years. This is due in part to the still limited, structural data available for these enzymes (<u>Amao, 2018</u>, <u>DeMirci, et al., 2022</u>). The complexity of the reactions that they catalyze, coupled to their diverse oligomeric forms, continues to pose challenges in deciphering their assembly, stability, and catalytic mechanisms. Addressing these knowledge gaps is essential to fully harnessing the knowledge and is critical for realizing the full biotechnological potential of ECRs in carbon capture and utilization (CCU) technologies. (<u>Bernhardsgrütter et al. 2021</u>)

In this work, we specifically focused on the ECR from *M. metallidurans*, a bacterium known for its resilience in metal-contaminated environments (<u>Diels et al. 2009</u>). This bacterium can thrive in the presence of heavy metals such as zinc, cadmium, and lead (<u>Von Rozycki et al. 2008</u>, <u>Diels et al. 2009</u>, <u>Janssen PJ et al. 2010</u>). Its enzymes, including ECRs, have developed unique adaptations to maintain function under environmental stress conditions (<u>Schada von Borzyskowski et al., 2013</u>). These make *M. metallidurans* an intriguing model system for studying the structural and functional properties of ECRs under extreme stress conditions, potentially providing valuable insights into enzyme stability and robustness.

An important aspect of ECRs is their oligomeric state, which is often linked to their stability and catalytic efficiency. Many ECRs, including those from organisms such as (*K. setae*), are known to form tetramers, which are thought to contribute to their enhanced catalytic function (<u>Erb et al., 2009</u>, <u>DeMirci, et al.,</u> <u>2022</u>). In contrast, our study identifies a novel dimeric form of ECR in *M. metallidurans*, marking a significant deviation from the typical tetrameric structure. This discovery suggests that dimerization versus tetramerization could play a crucial role in modulating enzyme activity and stability, possibly reflecting different evolutionary pressures or environmental adaptations (<u>Erb & Zarzycki, 2016</u>). The oligomeric state can have profound implications for enzyme function, influencing factors such as substrate binding affinity, reaction kinetics, and overall structural integrity (<u>Traut et al. 1994</u>). Tetrameric forms often exhibit cooperative binding (<u>DeMirci, et al., 2022</u>), where the interaction between subunits can enhance enzymatic efficiency, while dimeric forms may provide advantages in terms of flexibility and adaptability under specific environmental conditions (<u>Littlechild et al. 2015</u>). Understanding how these oligomeric states affect enzyme function is crucial for both fundamental enzymology and practical applications. This is particularly relevant in enzyme engineering efforts aimed at optimizing industrial CO_2_ fixation processes, as exemplified by studies on *B. ambifaria* (<u>Walker et al. 2015</u>). We collected X-ray crystallographic data at the Stanford Synchrotron Radiation Lightsource (SSRL), allowing us to resolve the intricate structural features of the *M. metallidurans* ECR dimer at an atomic resolution. The structural data provide valuable insights into how specific residues and inter-subunit interactions contribute to the stability of the dimeric form. Additionally, *in-silico* modeling allowed us to hypothesize how these residues might prevent tetramer formation, potentially by destabilizing the dimer-dimer interface <u>(Goldenzweig et al. 2018</u>). By comparing these findings with the tetrameric ECR structure of *K. setae*, we aimed to delineate the structural basis of their different oligomeric preferences and the functional implications of these differences.

In addition to the structural studies, we also conducted a comparative analysis of the primary sequences of ECRs from *M. metallidurans* and *K. setae*, focusing on regions associated with secondary structural elements, such as alpha helices and beta strands (<u>Todd et al. 2001</u>). This analysis helped identify specific sequence adaptations that may underpin the observed differences in oligomeric state. These sequence variations likely represent evolutionary responses to the distinct ecological niches occupied by these organisms, affecting enzyme stability and activity under different environmental conditions.

The identification of a dimeric ECR structure in *M. metallidurans* represents an important step forward in understanding the structural diversity of CO_2_-fixing enzymes. This discovery challenges the conventional notion that tetrameric forms are essential for catalytic efficiency in ECRs and opens new avenues for exploring how dimerization might confer unique advantages in terms of enzyme stability and adaptability (<u>Mandeep et al. 2020</u>). By elucidating the structural and functional differences between the dimeric and tetrameric forms, we aim to provide a deeper understanding of the relationship between enzyme structure, function, and adaptation, which is crucial for advancing enzyme engineering efforts aimed at enhancing carbon sequestration capabilities.

The overarching goal of this work is to contribute to the growing body of knowledge on CO_2_ fixation and to pave the way for the development of more efficient catalytic processes for carbon capture and conversion. The insights gained from studying the structural basis of ECR oligomerization not only advance our understanding of these enzymes but also hold potential for their applications in biotechnological and industrial processes aimed at reducing atmospheric CO_2_ levels, there by addressing pressing environmental challenges (<u>Gupta CL et al. 2014</u>).

## 2. Results

### 2.1. *M. metallidurans* ECR forms dimer in solution rather than canonical tetramer

In this study, we explored the oligomeric states of Enoyl-CoA carboxylases/reductases (ECRs) from *M.metallidurans* and *B. ambifaria*, focusing on their structural and biochemical distinctions. Specifically, the dimeric form of the ECR from *M. metallidurans* ECR^Mm_Dim^ was examined in comparison with the canonical tetrameric structure characteristic of *B. ambifaria* ECR^Ba_Tet^. To distinguish these oligomeric states, we employed size exclusion chromatography (SEC), which revealed clear differences in their elution profiles (Figure 1). The ECR^Ba_Tet^ eluted earlier, reflecting its larger molecular weight, while the ECR^Mm_Dim^ exhibited a later elution, consistent with its smaller molecular size. Supplementary Figure 1 provides a comprehensive set of chromatograms illustrating the behavior of these proteins in SEC, both as individual samples and in mixed populations. Notably, the unique elution profiles highlight fundamental differences in their assembly mechanisms and potential interaction properties, which may underlie the distinct functional roles of these enzymes. Co-elution experiments involving a combination of both proteins revealed complex interaction patterns, further underscoring the divergent assembly and stability characteristics of their oligomeric states.

**Figure 1:**
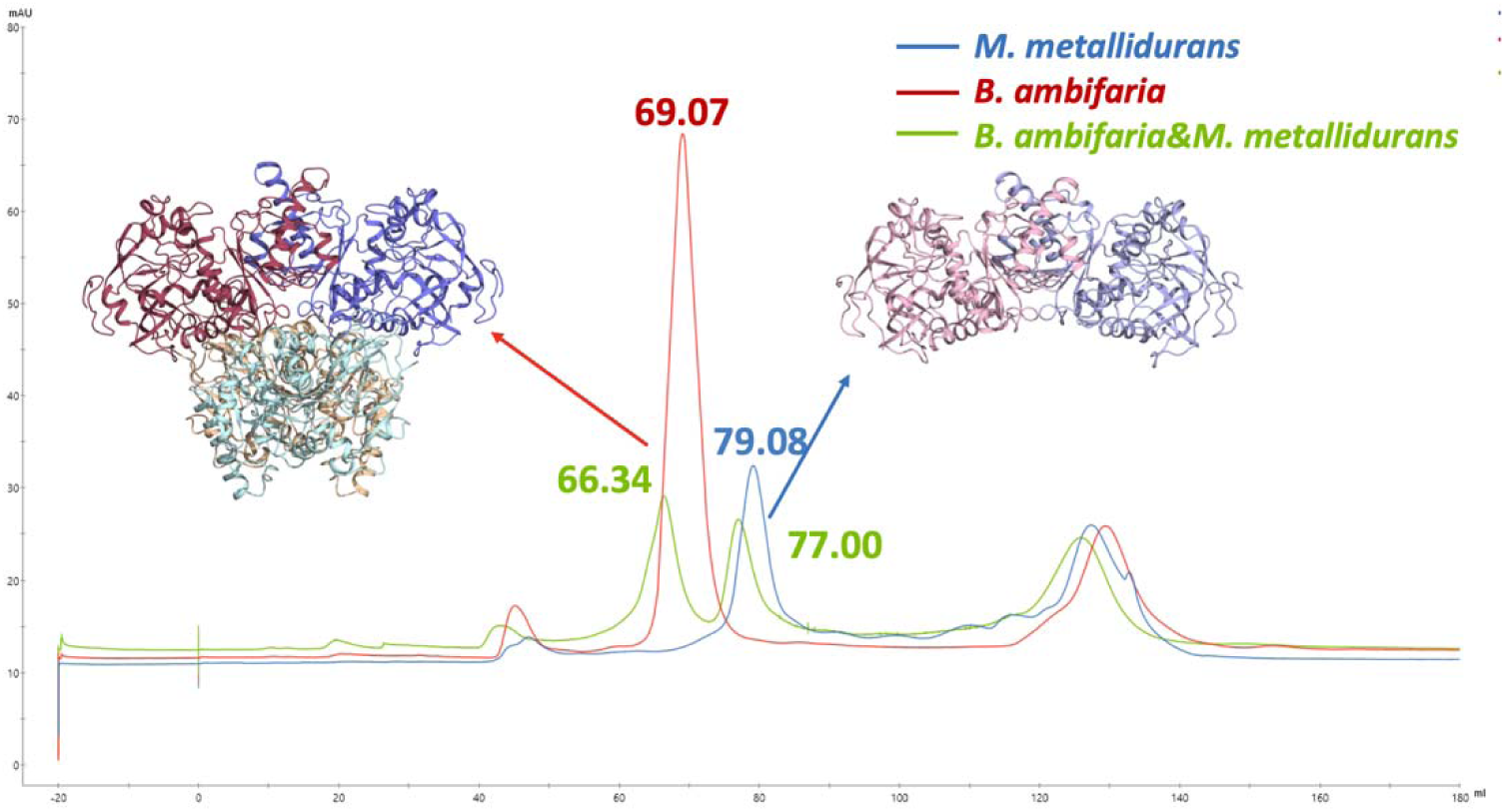
Comparative analysis of size exclusion chromatography for dimeric protein *M. metallidurans* and tetrameric protein *B. ambifaria*, this figure compares the size exclusion chromatography (SEC) profiles of the tetrameric protein *B. ambifaria* and the dimeric protein *M. metallidurans*. The blue chromatogram represents the profile of pure *M. metallidurans* protein, while the red chromatogram corresponds to the 1 mL SEC profile of pure *B. ambifaria* protein at the same elution volume. The green chromatogram represents a mixture of equal volumes (500 µL each) of *B. ambifaria* and *M. metallidurans* proteins.

Before delving into the detailed structural analysis of the ECR^Mm_Dim^, we present the results of its crystal structure determination to establish a foundational understanding of its overall architecture. The ECR^Mm_Dim^ adopts a distinctive structural arrangement that diverges significantly from the canonical tetramer, as observed in related systems. The crystal structure highlights key features, including the organization of its active site residues and the spatial arrangement of regions critical for dimer stabilization. To gain deeper insights into its dynamic properties, we conducted a thorough atomic-level motion analysis using B-factors derived from the crystallographic data. This was further complemented by ellipsoidal representations, providing a visual depiction of regions with differential flexibility and structural stability. These analyses revealed notable trends in the relative rigidity and mobility of specific domains within the dimer, offering clues about their potential influence on enzymatic activity and functional efficiency.

Collectively, these findings set the stage for a detailed examination of how the oligomeric state of the ECR^Mm_Dim^ shapes its function, interactions, and assembly mechanisms, compared to its tetrameric ECR^Ba_Tet^ counterpart.

### 2.2. Structural Analysis of ECR^Mm_Dim^

Before delving into the detailed structural analysis of the ECR^Mm_Dim^, we first present the fundamental results of its crystal structure determination and refinement (Table 1). The crystal structure was solved in the space group C 1 2 1, with unit cell dimensions of *a* = 187.99 Å, *b* = 104.31 Å, *c* = 44.81 Å, and angles α = 90°, β = 91.6°, γ = 90°. Data was collected at a synchrotron source at 100 K, with a wavelength of 0.980 Å. The dataset was refined to a high resolution of 1.40 Å, achieving 98.0% completeness (85.5% in the highest resolution shell), a redundancy of 6.83 (4.98), and a CC(1/2) of 1.0 (0.66). The structural coordinates have been deposited in the PDB under accession ID 9KVA (Table 1).

**Table 1:** Data collection and refinement statistics.

| **Data Collection** |  |
| --- | --- |
| **Space group** | C 1 2 1 |
| **Cell Dimensions** |  |
| **a, b, c (Å)** | 187.99 104.31 44.81 |
| **α, β, γ (°)** | 90.0 91.6 90.0 |
| **Resolution (Å)** | 35.36-1.40 (1.43-1.40) |
| **Rsym or Rmerge** |  |
| **Rmeas** | 6.7(165.7) |
| **I / σI** | 13.63 (1.46) |
| **CC(1/2)** | 1.0 (0.66) |
| **Completeness (%)** | 98.0 (85.5) |
| **Redundancy** | 6.83 (4.98) |
| **Wavelength (Å)** | 0.980 |
| **Temperature** | 100 K |
| **Data collection** | Synchrotron |
| **Oligomer** | Homodimer |
| **Refinement** |  |
| **Resolution (Å)** | 35.4-1.40 (1.43-1.40) |
| **No. reflections** | 158404 (10156) |
| **Rwork / Rfree** | 0.15/0.18 (0.32/0.34) |
| **No. atoms** |  |
| **Protein** | 6224 |
| **Ligand/ion** | 32 |
| **Water** | 599 |
| **B-factors** |  |
| **Protein** | 33.72 |
| **Ligand/ion** | 40.02 |
| **Water** | 42.43 |
| **R.m.s. deviations** |  |
| **Bond lengths (Å)** | 0.009 |
| **Bond angles (°)** | 1.692 |
| **Ramachandran Statistics (%)** |  |
| **Favored** | 93.9% |
| **Allowed** | 6.1% |
| **Data Collections** |  |
| **Space group** | C 1 2 2 |
| **Cell Dimensions** |  |
| **a, b, c ( Å)** | 187.99 104.31 44.81 |
| **α, β, γ (°)** | 90.0 91.6 90.0 |
| **Outliers** | 0 |
| **PDB ID** | 9KVA |

The crystallographic refinement yielded *R_work* and *R_free* values of 0.15 and 0.18, respectively, with no Ramachandran outliers and 93.9% of residues in favored regions. The structure consists of 6224 protein atoms, 32 ligand/ion atoms, and 599 water molecules, with B-factors averaging 33.72 Å² for the protein, 40.02 Å² for ligands, and 42.43 Å² for water molecules. Root means square (R.m.s.) deviations for bond lengths and angles were 0.009 Å and 1.692°, respectively, indicating high model accuracy.

The asymmetric unit comprises a homodimer of ECR^Mm_Dim^, with Chains A and B forming a near-symmetric arrangement (Figure 2A). This symmetry is retained upon rotation, showcasing the inter-chain interaction regions and the back face of the dimer (Figure 2B). These foundational results provide critical context for interpreting the dimer’s dynamic properties and its role in catalytic functions. A detailed structural analysis of the ECR^Mm_Dim^ was conducted, focusing on atomic-level motions through B-factor analysis and ellipsoidal representations of the protein’s structural stability. This study revealed regions of both flexibility and rigidity, identifying potential hotspots for functional conformational changes (Figure 2). The overall arrangement and symmetry between Chain A (light pink) and Chain B (light blue) are illustrated in a cartoon representation of the dimer, viewed from the front face (Figure 2A). When the ECR^Mm_Dim^ is rotated 180°, the back face of the dimer is revealed, offering an alternative perspective to better understand the interaction regions and the spatial relationship between the two chains (Figure 2B). Next, the electrostatic surface potential of the ECR^Mm_Dim^ is analyzed. The front face shows charge distribution, with red representing a negative potential (-5 kT/e) and blue representing a positive potential (+5 kT/e), revealing areas that may participate in protein-protein interactions or ligand binding (Figure 2C). The back face of the ECR^Mm_Dim^’s electrostatic potential is shown for comparison, highlighting contrasting electrostatic properties that could influence binding or interaction regions (Figure 2D). B-factor analysis of the solvent-accessible surface is displayed for the front face of the dimer, where regions with higher B-factors, colored yellow-red, indicate more flexible areas, and blue-green regions represent more rigid areas (Figure 2E). These flexible regions could be involved in substrate binding and catalytic activity. The same analysis for the back face further complements the understanding of flexibility and rigidit across both face (Figure 2F).

**Figure 2:**
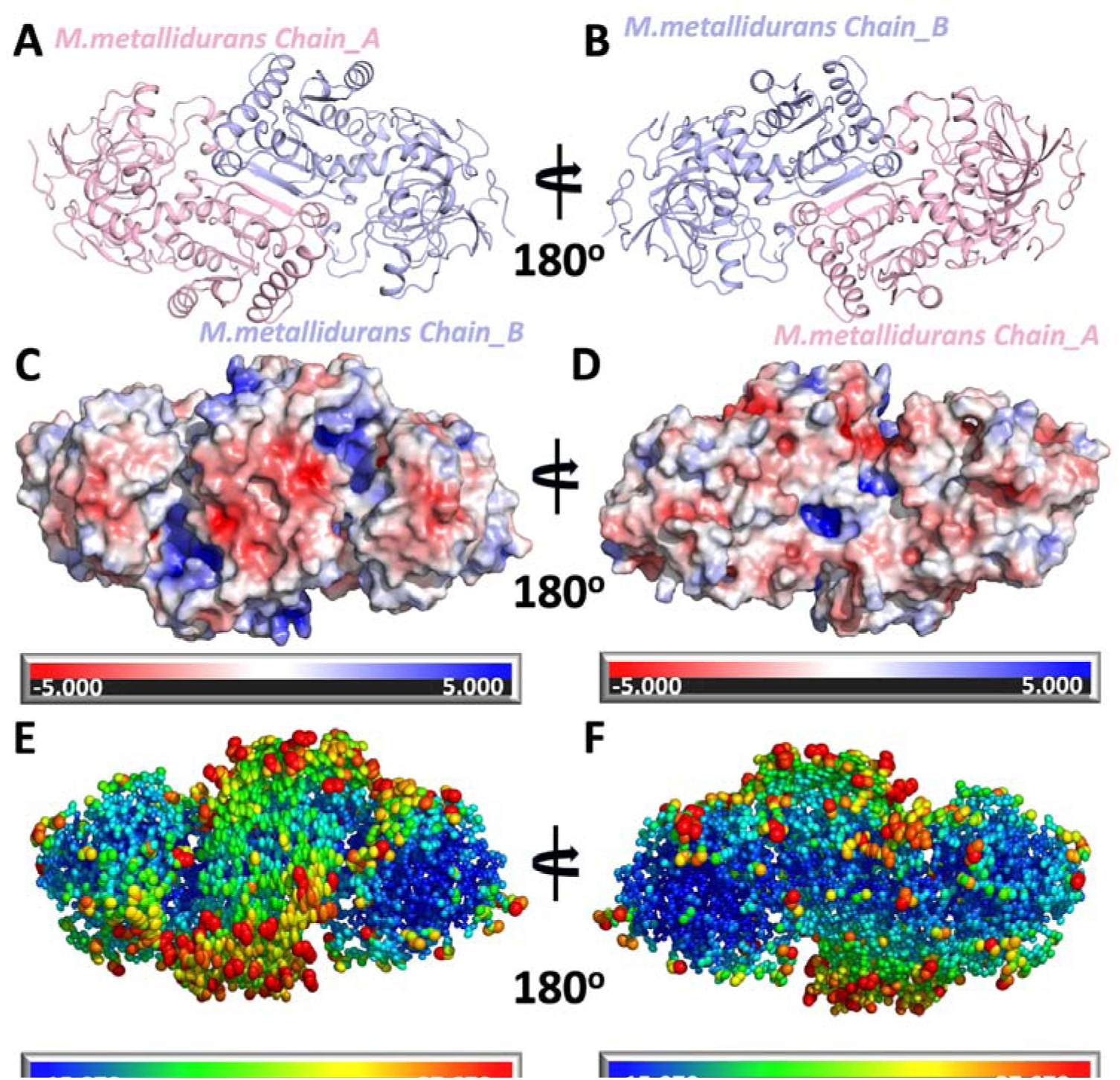
Structural, electrostatic surface, and B-factor representations of *M. metallidurans*. **a)** Cartoon representation of the dimeric structure of *M. metallidurans*, with Chain A shown in light pink and Chain B in light blue. This view represents the front face of the dimer. **b)** The structure shown in panel (A) is rotated to show the back face of the dimer. **c)** Electrostatic surface potential representation of the front face of the *M. metallidurans* dimer. The surface is colored according to electrostatic potential, with red representing negative potential (-5 kT/e) and blue representing positive potential (+5 kT/e). **d)** The electrostatic surface potential of the back face of the dimer, corresponding to the view in panel (B). **e)** Surface representation of the front face of the *M. metallidurans* dimer, colored according to B-factors and ellipsoidal shapes according to carbon alpha. **f)** The surface representation of the back face of the dimer, corresponding to the view in panel (B). **Note:** Panels A, C, and E represent the front face of the dimer, while panels B, D, and F represent the back face. The views are rotated 180° relative to each other to illustrate different perspectives.

This structural analysis suggests that the ECR^Mm_Dim^ exhibits specific folding and dynamic behaviors that are critical to its function and stability. Supplementary Figure 6, which includes AlphaFold predictions, supports the comparison with potential tetrameric forms.

### 2.3. Role of Crystal Packing in Modulating Enzymatic Behavior

The crystallographic analysis of ECR^Mm_Dim^ revealed a unique dimeric arrangement within the crystal lattice, distinguishing it from the previously characterized tetrameric structures of enoyl-CoA carboxylase/reductase (ECR) enzymes from other species (Figure 3). Crystal packing does not influence the selection of dimeric or tetrameric forms.

**Figure 3:**
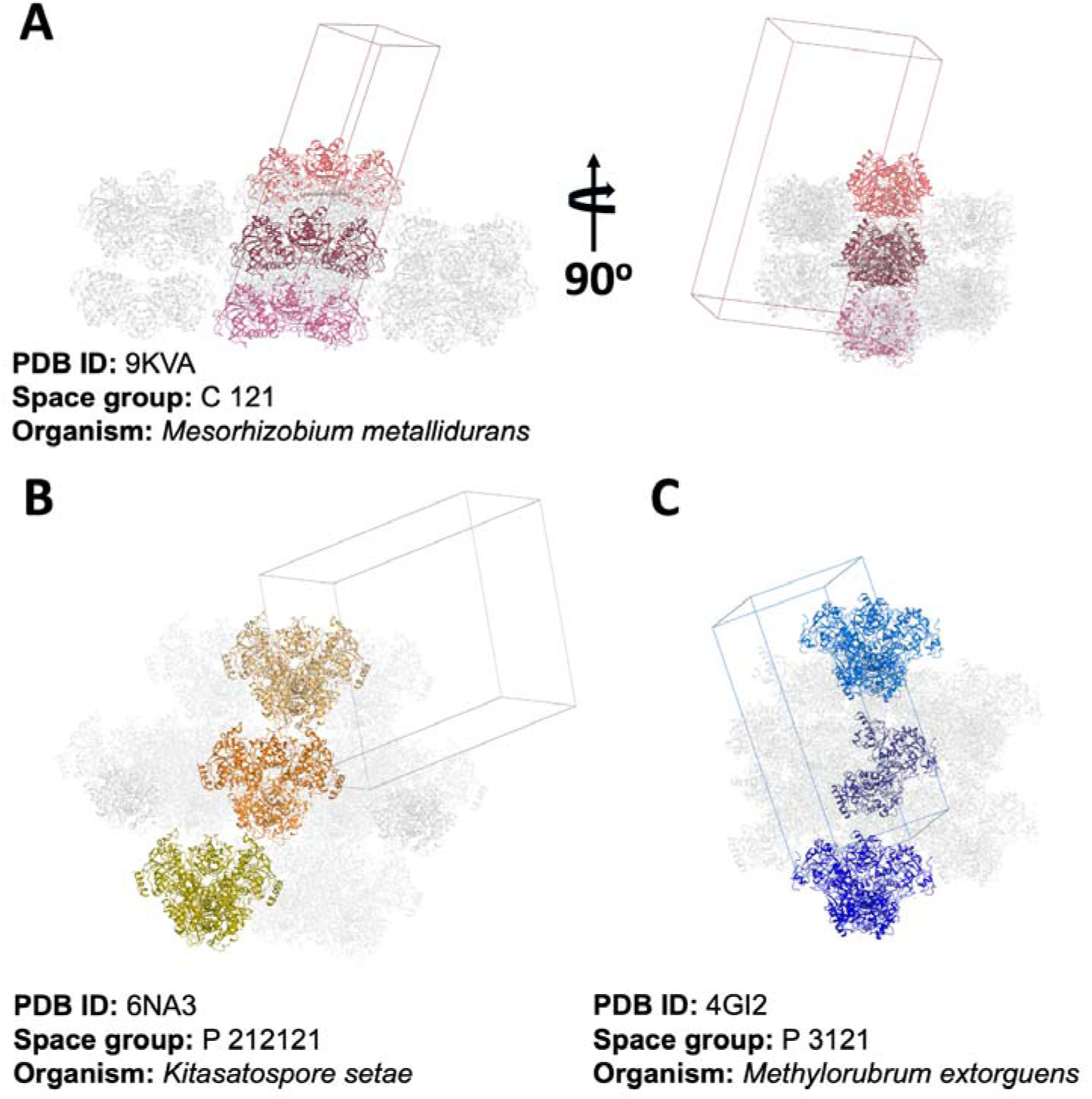
Crystal packing of ECR structures from different organisms. **a)** *M. metallidurans* ECR (PDB ID: 9KVA; space group C 1 2 1) showing a dimeric assembly in two orthogonal orientations, highlighting subunit arrangement and interface. **b)** *K. setae* ECR (PDB ID: 6NA3; space group P 21 21 21) with a canonical tetrameric assembly, distinct from the dimeric form in *M. metallidurans*. **c**) *M. extorquens* ECR (PDB ID: 4GI2; space group P 31 2 1) displaying tetrameric packing despite being annotated as a dimer, indicating possible misclassification or structural plasticity.

A detailed comparison with the tetrameric ECR structures from *Kitasatospora setae* (PDB ID: 6NA3) (ECR^Ks_Tet^) and *Methylorubrum extorquens* (PDB ID: 4GI2)(ECR^Me_Tet^) showed marked differences in crystal packing patterns. These differences include the orientation and positioning of subunits within the asymmetric unit, the nature of inter-subunit contacts, and overall inter-oligomeric interactions within the lattice. In ECR^Mm_Dim^, the dimers are stabilized through contacts specific to their crystallographic arrangement, which are absent or reconfigured in the tetrameric structures of other species.

The dimeric and tetrameric forms of ECR enzymes represent intrinsic structural and functional states, with crystal packing serving as a stabilizing factor for the oligomer present under the crystallization conditions. In contrast to tetrameric ECRs, where inter-dimer interfaces are essential for enzymatic function, the ECR^Mm_Dim^ structure maintains its stability and activity through intra-dimer interactions that are independent of crystal packing constraints.

Supplementary Figure 3 illustrates differences in electrostatic surface potentials between dimeric and tetrameric structures. The surface charge distribution of the dimeric structure contributes to its function, but these electrostatic patterns arise from the intrinsic architectural differences between the dimeric and tetrameric forms, rather than crystal packing. These differences may influence substrate or protein interactions but do not dictate oligomeric preferences.

The crystal packing of ECR^Mm_Dim^ is species-specific and does not determine the enzyme’s quaternar structure. The dimeric arrangement, stabilized by intra-dimer interactions, remains functionally distinct and separate from the tetrameric assemblies observed in other ECRs.

### 2.4. Residues Preventing Tetramer Formation in *M. metallidurans*: Structural and *In-Silico* Insights

Through a combination of structural analysis and in-silico modeling, specific residues in *M. metallidurans* were identified as key inhibitors of tetramer formation (Figure 4). Notably, Glu224, Ala225, Gly226, and Gly227 were found to play critical roles in destabilizing the interaction surfaces necessary for tetramerization, thus maintaining the enzyme in its dimeric state. In the sequence alignment between *M. metallidurans* and *K. setae*, key residues involved in the dimer interface are boxed, emphasizing their structural significance (Figure 4a). The overall interaction surface stabilizing the dimer is displayed in the ECR^Mm_Dim^, where chain A (pink) and chain B (blue) are shown interacting (Figure 4b). A close-up view of chain A residues (Asp129, Pro128, Ala127, Val131, Ser132, Val130, and Val133) reveals their tight packing within the interface, which is crucial for dimer stability and prevents tetramer formation (Figure 4c). Additionally,residues Pro47, Gly48, and Glu49 from chain B contribute to this stability by interacting Asp129, Ala127, and Pro128 at the core of the dimer interface (Figure 4d). Chain B residues Val383, Leu362, and Asp361 may further stabilize the dimer through hydrophobic and electrostatic interactions with chain A, as depicted in the surface representation (Figure 4e). Chain A residues (Asp129, Ala127, and Pro128) are positioned to maintain the integrity of the dimer interface, highlighting their role in ensuring the structure remains dimeric (Figure 4f). Finally, the interaction between chain B residues (Ala225, Gly226, and Gly227) is shown to contribute to the stabilization of the dimer interface and reinforces the exclusion of tetramer formation (Figure 4g).

**Figure 4:**
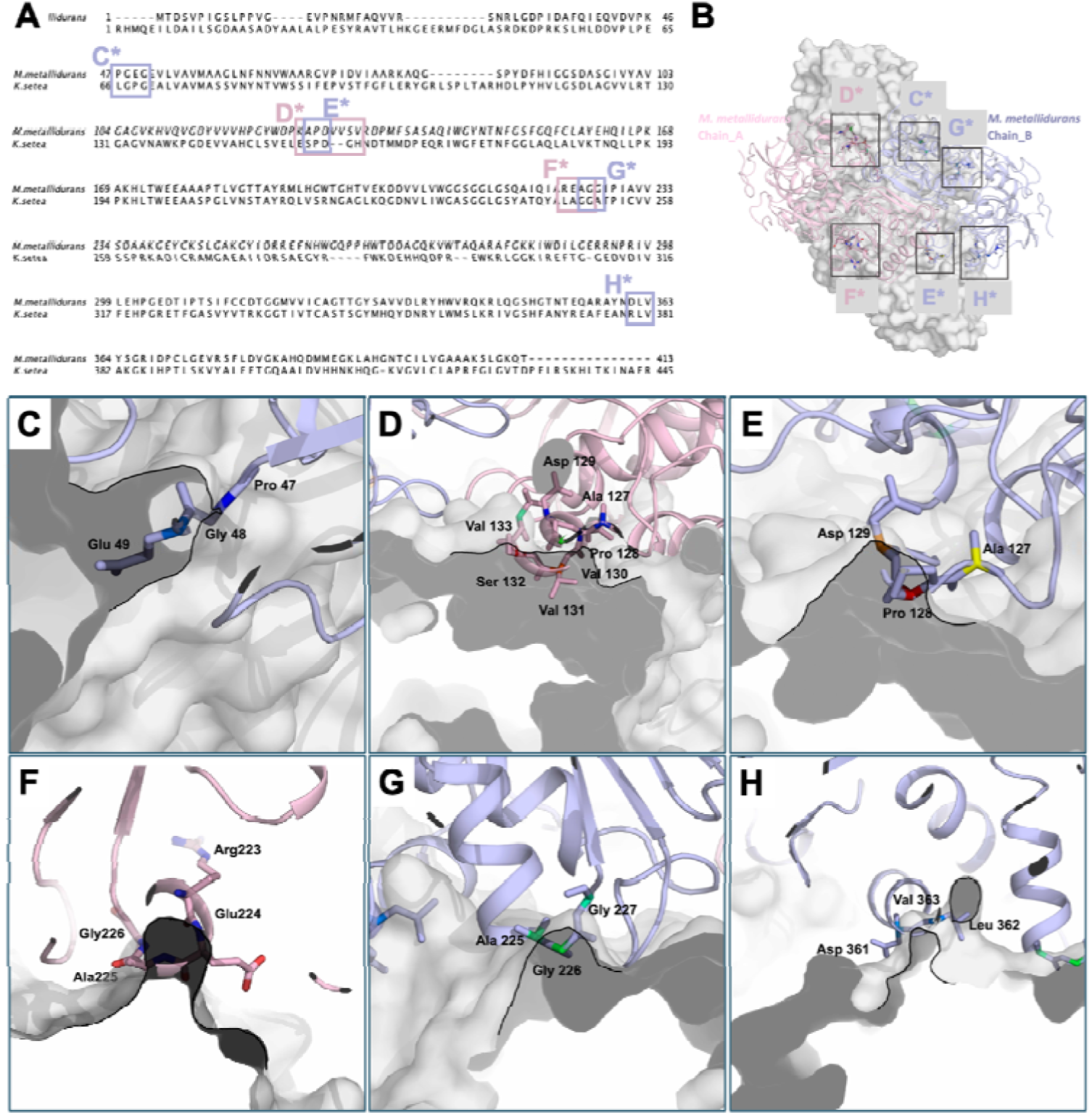
Structural analysis of the ECR dimer from *M. metallidurans*. **a)** Sequence alignment of *M. metallidurans* and *K. setae*, highlighting crucial residues at the dimer interface. These residues are boxed to underscore their structural and functional importance in forming tetramers in *M*. *metallidurans*. Details of the regions important in the dimer interface of the *M. metallidurans* dimer are shown in panels B-G, displaying chains A (pink) and B (blue), with the surface of each chain. **b)** Overview of the dimeric structure of *M. metallidurans* ECR. Chain A is shown in pink and Chain B in blue, with key regions at the dimerization interface highlighted as boxed areas (C*-H*). These boxes correspond to the close-up views presented in panels C-H, emphasizing the critical residues involved in dimer stabilization. **c)** Close-up view of chain B residues (Pro47, Gly48, and Glu49) at the dimer interface. These residues form a key region contributing to the stability of the dimerization surface in *M. metallidurans*. Pro47 introduces a sharp bend, while Gly48 provides structural flexibility, and Glu49 plays a potential role in electrostatic interactions. Together, these residues define the conformation necessary for maintaining proper dimer assembly. **d)** Close-up of key interface residues in chain A (Asp129, Pro128, Ala127, Val131, Ser132, Val130, and Val133). **e)** In-depth look at chain B residues (Pro47, Gly48, and Glu49) at the dimer interaction surface. **f)** Surface representation of chain B residues (Val383, Leu362, and Asp361). **g)** Analysis of chain A residues (Asp129, Ala127, and Pro128) at the interface. **h)** Interaction of chain B residues (Ala225, Gly226, and Gly227) at the interface.

These findings were further validated by *in-silico* models, which provided detailed visualizations of the regions preventing tetramer assembly. The study also included the in-silico construction of a potential tetrameric *M. metallidurans* ECR (ECR^Mm_Tet^), enabling a direct comparison with naturally occurring tetramers from other species (Supplementary Figure 2). These models revealed significant differences in stabilization mechanisms, particularly at the tetramer interfaces. The ECR^Mm_Dim^ may have evolved to avoid tetramerization due to environmental or functional pressures. Collectively, these insights offer a molecular explanation for the distinct oligomeric states of *M. metallidurans* compared to other species, such as *B. ambifaria* and *K. setae*, and provide a hypothetical framework for understanding the evolutionary adaptations of these enzymes under different conditions.

Expanding on the structural analysis of ECR^Mm_Dim^, a comparative study with ECR^Ks_Tet^ reveals critical differences in their oligomerization behaviors. While ECR^Mm_Dim^ is stabilized in a dimeric form by residues such as Glu224, Ala225, Gly226, and Gly227, these residues prevent the interaction surfaces required for tetramerization. In contrast, *K. setae* readily forms a tetramer, with the interface between Chains A, B, C, and D stabilized by additional contacts, particularly through hydrophobic and electrostatic interactions (Figure 5). In ECR^Ks_Tet^, specific residues like Asp129, Val130, and Pro128 in Chain A engage in key interactions with residues in adjacent chains, reinforcing the tetrameric structure (Figure 5B). These residues play a significant role in bridging the interface between Chains A and C, as well as Chains B and D, promoting the assembly of a stable tetramer. On the other hand, in ECR^Mm_Dim^, the corresponding residues in Chain A are arranged in a way that disrupts such contacts, effectively preventing tetramer formation (Figure 5C). Notably, the positioning of Glu224 and Ala225 at the dimer interface acts as a barrier to further oligomerization, maintaining the enzyme in its dimeric form.

**Figure 5:**
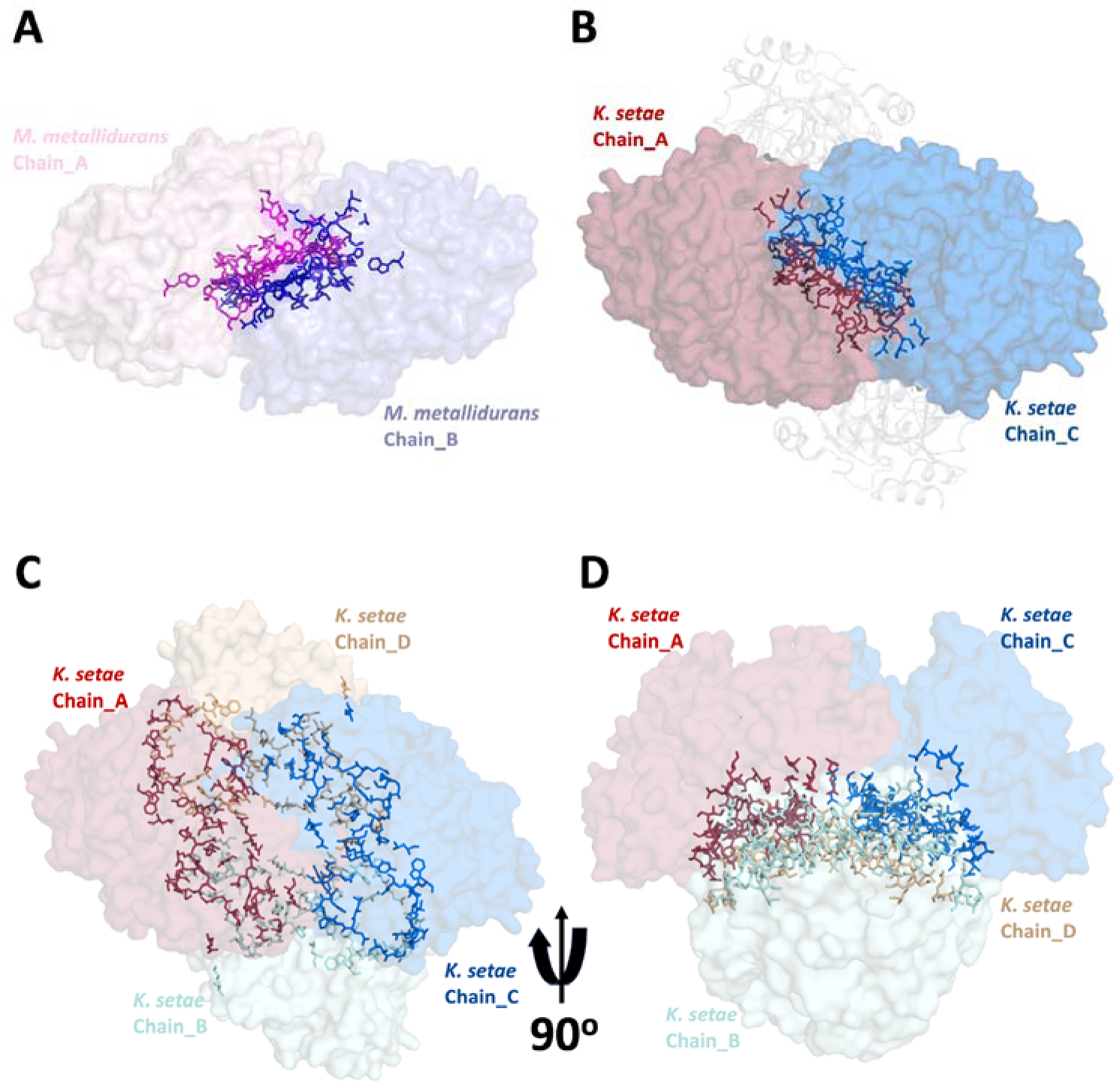
Comparison of the dimer and tetramer interfaces of *M. metallidurans* and *K. setae* ECRs. **a)** The surface representation of the *M. metallidurans* ECR dimer, shows Chain A and Chain B. The dimerization interface is highlighted, with interacting residues displayed as sticks. **b)**The *K. setae* ECR dimer structure, showing the dimerization of Chain A and Chain C. The interaction interface between these two chains is highlighted, with the interacting residues displayed as sticks**. c)** Tetrameric structure of *K. setae* ECR, showing Chains A, B, C, and D. The tetramer interface between the chains is shown, with interacting residues displayed as sticks. **d)** A 90° rotated view of the *K. setae* tetramer further illustrates the chains’ spatial arrangement and the extended tetramerization interface.

Through *in silico* modeling and structural comparison, it becomes evident that *M. metallidurans* has evolved to favor a dimeric configuration, likely as a response to specific functional or environmental pressures. By contrast, *K. setae*’s tetrameric structure appears to confer stability and function under different conditions. These insights not only elucidate the molecular mechanisms behind the distinct oligomeric states of these species but also offer a broader understanding of how such enzymes may adapt to varying environmental or metabolic needs.

### 2.5. Comparative Study of Primary Sequences

A comparative analysis of the primary sequences of *M. metallidurans* and *K. setae* ECRs was conducted, focusing on regions associated with secondary structural elements like helices and beta strands (Figure 6). Significant variations were observed in these regions, particularly in the loops connecting ke structural domains. These differences were mapped and categorized into regions K1-K5 for *K. setae* and M1-M4 for *M. metallidurans*, offering a structural basis for their divergent functional properties. Supplementary Figure 4 illustrates these structural elements and their differences, providing a structural foundation for understanding the functional adaptations of both enzymes.

**Figure 6:**
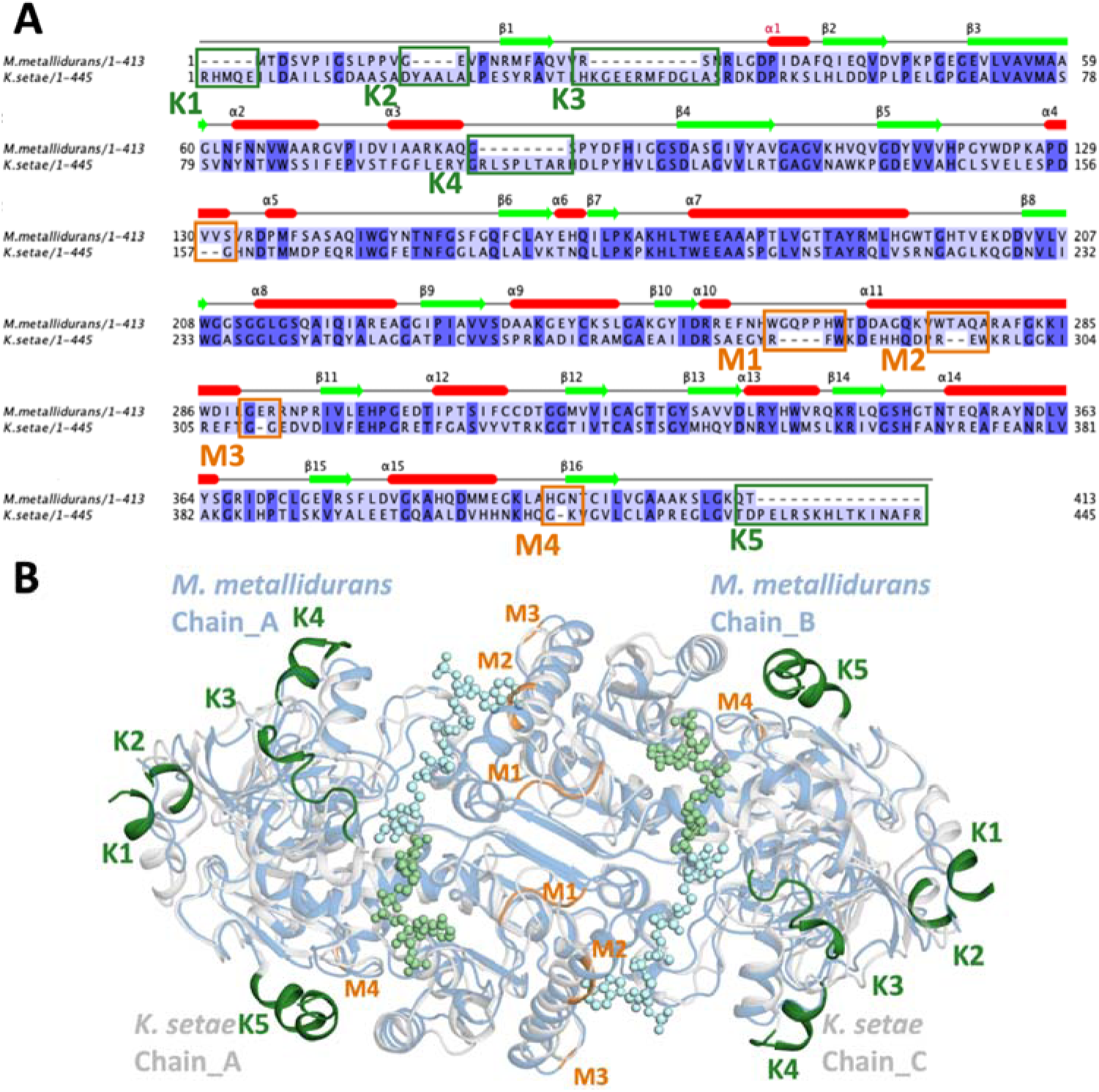
Comparative sequence and structural analysis of ECRs from *M. metallidurans* and *K. setae*, **a)** Sequence alignment of ECRs from *M. metallidurans* (1–413) and *K. setae* (1–445). The secondary structural elements are annotated above the sequences, with helices represented as red cylinders and beta strands as green arrows. Key regions that differ between the two sequences are highlighted. Differences in the *M. metallidurans* structure compared to *K. setae* are labeled as K1-K5 (green), while differences in *K. setae* compared to *M. metallidurans* are labeled as M1-M4 (orange). **b)** Structural superposition of the *M. metallidurans* dimer (cyan) and the open subunits of the apo *K. setae* tetramer (blue-white). Regions that differ in the *M. metallidurans* structure compared to *K. setae* are highlighted and labeled as K1-K5 (green), while regions that differ in *K. setae* compared to *M. metallidurans* are labeled as M1-M4 (orange). These variations are shown to impact the overall structural differences between the two ECRs.

### 2.6. Structural and Functional Divergence in ECR NADPH binding sites

A comparative structural analysis of the NADPH binding sites in ECR^Mm_Dim^ and ECR^Ks_Tet^ was performed to explore the molecular basis of their distinct catalytic properties. Figure 7 provides insights into the apo and NADPH-bound conformations of both enzymes, highlighting differences in their active site architectures and ligand-induced conformational changes. In ECR^Mm_Dim^, the apo structure adopts a relatively open conformation, with key active site residues such as Ala79 and Arg279 positioned 21.1 Å apart, allowing for solvent accessibility and a stable active site environment (Figure 7A). Upon NADPH binding, a subtle yet significant rearrangement occurs, where Ala79 shifts inward by 1.8 Å, narrowing the active site entrance (Figure 7B). This conformational shift suggests a modest degree of induced fit, though with limited flexibility in substrate accommodation.

**Figure 7:**
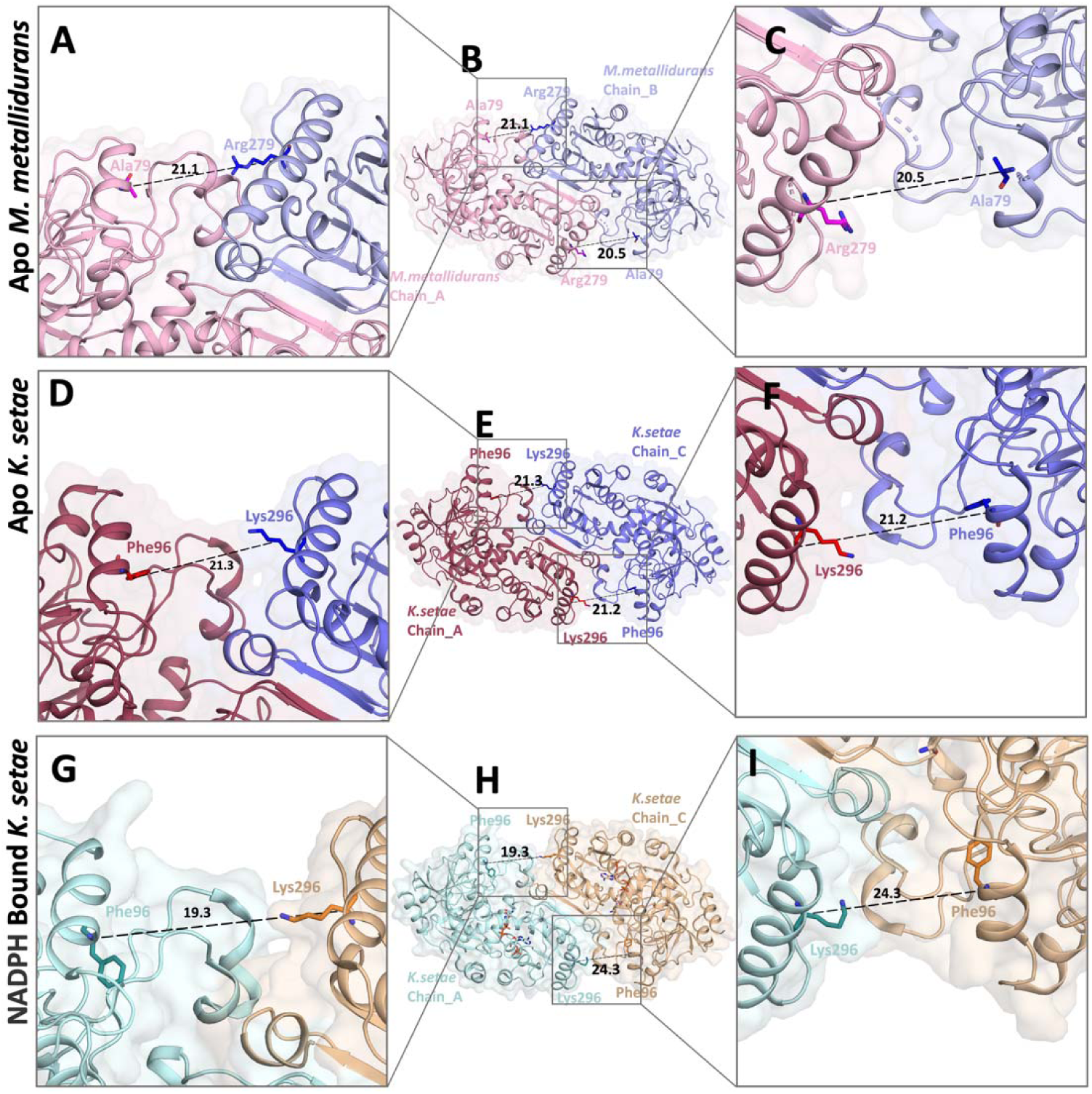
Comparison of Substrate Binding Regions, **a, b, c)**These panels show differences between the substrate binding regions in the open A-B subunits of the apo *M. metallidurans* structure. **d, e, f)** These panels depict differences between the substrate binding regions in the open A-C subunits of the apo *K. setae* structure (PDB ID: 6NA3). **g, h, i)** These panels highlight differences in the substrate binding regions between the open A-C subunits of the NADPH-bound *K. setae* structure (PDB ID: 6NA6).

In contrast, ECR^Ks_Tet^ exhibits a more dynamic active site arrangement. In the apo form, the active site residues Phe96 and Lys296 are separated by 24.3 Å, maintaining a slightly expanded pocket (Figure 7D). Upon NADPH binding, a pronounced conformational change occurs, bringing Phe96 and Lys296 closer together to 19.3 Å, which significantly alters the topology of the active site (Figure 7E). This structural reorganization likely enhances NADPH binding efficiency and optimizes substrate positioning, thereby improving catalytic performance. The superimposition of the apo and NADPH-bound structures of both enzymes (Figure 7G) further underscores these differences, emphasizing how the tetrameric nature of ECR^Ks_Tet^ promotes greater conformational adaptability compared to the more rigid dimeric structure of ECR^Mm_Dim^.

A detailed comparative analysis of the active sites inECR^Mm_Dim^ and ECR^Ks_Tet^ revealed key differences in the residues surrounding the NADPH binding site (Figure 8). Notably, ECR^Ks_Tet^ possesses unique features in residues such as Pro120, Ile144, and Phe366, which contribute to improved substrate positioning and catalytic efficiency (Figure 8A). These residues engage in hydrophobic and electrostatic interactions, shaping an active site architecture that facilitates efficient NADPH binding and stabilization. Additionally, the tetrameric interface provides further stabilization through inter-subunit contacts, reinforcing active site accessibility (Figure 8B).

**Figure 8:**
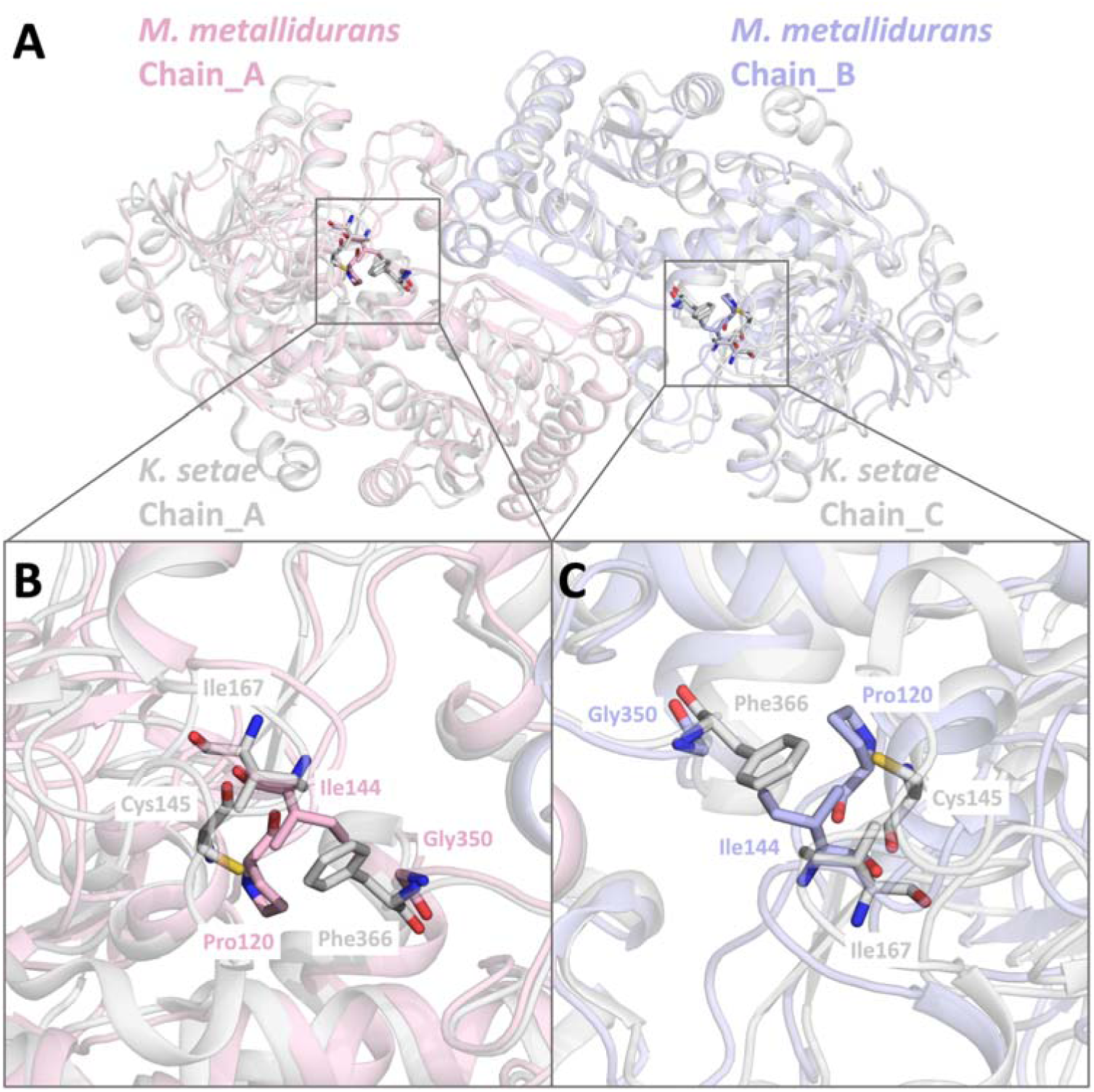
Comparison of active site differences between *M. metallidurans* and *K. setae* ECR structures, **a)** Conformational changes in the active site residues of the dimeric *M. metallidurans* structure are aligned with the A-C open subunit of the apo form *K. setae* tetramer structure. The alignment highlights the structural differences and similarities in the active sites. **b)** Detailed view of the active site residues between *M. metallidurans* chain A (light pink) and *K. setae* chain A (raspberry). Key residues and the NADPH molecule are labeled, showing the positions and interactions within the active site. The residues shown include Ile167, Ile144, Cys145, Gly350, and Phe366. **c)** Detailed view of the active site residues between *M. metallidurans* chain B (light blue) and *K. setae* chain C (marine). Key residues and the NADPH molecule are labeled, illustrating the conformational changes. Residues shown include Ile167, Ile144, Cys145, Gly350, and Phe366.

Conversely, the dimeric structure of ECR^Mm_Dim^ presents a more restricted active site conformation. Residues such as Cys145 and Gly350 are positioned in a manner that may limit substrate dynamics (Figure 8D). While this arrangement could enhance stability under specific environmental conditions, it might also impose constraints on catalytic efficiency. The spatial organization of these residues (Figure 8E) supports the hypothesis that ECR^Mm_Dim^ has evolved to prioritize stability over catalytic flexibility.

The oligomeric state plays a crucial role in determining the catalytic mechanisms and efficiency of these enzymes. The structural organization observed in ECR^Mm_Dim^ and ECR^Ks_Tet^ provides a molecular explanation for the distinct functional properties of dimeric and tetrameric forms. The tetrameric assembly of ECR^Ks_Tet^ enables enhanced NADPH accessibility through a network of inter-subunit interactions, whereas the dimeric nature of ECR^Mm_Dim^ imposes spatial constraints on substrate binding. These structural insights, reinforced by AlphaFold-predicted models (Supplementary Figure 7), offer a deeper understanding of the functional adaptations of these enzymes. The interplay between active site architecture and oligomeric organization underscores their evolutionary divergence, with potential implications for enzyme engineering and biotechnological applications.

## 3. Discussion

The findings emphasize the necessity of investigating tetrameric and dimeric forms of enzymes to comprehensively understand their functional roles and adaptive mechanisms. The distinct SEC profiles for ECR^Mm_Dim^ and ECR^Ba_Tet^ proteins underscore how quaternary structure significantly impacts protein behavior and function. Previous research has shown that variations in oligomeric states can lead to substantial differences in catalytic activities, particularly in extremophilic organisms where structural conformation is closely linked to environmental adaptation (<u>Littlechild et al. 2015</u>).

These findings have broader implications for understanding the diverse strategies employed by microorganisms to adapt to extreme environmental conditions. Environmental adaptations can exert evolutionary pressure on enzyme structure-function relationships, driving the adaptation of quaternary structures across a wide variety of organisms.

The structural analysis of the ECR^Mm_Dim^, especially through ellipsoidal B-factor representations, provides critical information on the protein’s stability and folding mechanisms. Similar studies on protein dimers have highlighted their role in maintaining structural integrity under various environmental stresses <u>(Flick K et al. 2012</u>). Understanding atomic motion and rigidity within the dimeric structure is essential for advancing knowledge in protein chemistry. These insights are vital for applications aimed at enhancing protein stability for industrial use, where enzymes must tolerate thermal and mechanical stresses in various processes. These insights have potential applications in the design of proteins with enhanced stability and functional robustness under various conditions, which is valuable for both scientific and industrial purposes (<u>Sun et al. 2019</u>). The crystallographic examination of ECR^Mm_Dim^ provides valuable insights into how crystal packing influences enzyme activity. The arrangement of molecules within the crystal lattice plays a crucial role in enzyme function, especially in drug design, where the binding of inhibitors or other small molecules depends heavily on the accessibility and stability of the active site. This finding supports prior research demonstrating that crystal packing can modulate enzyme activity by affecting the structural configuration of the active site, making it either more or less accessible to small molecules (<u>Honarparvar et al. 2013; Goodford et al. 1984</u>). Understanding these stability characteristics is particularly important for future engineering efforts aimed at optimizing enzymes for industrial processes. Enzymes often encounter significant thermal and mechanical stresses during such processes, and the ability to design proteins that can withstand these conditions is crucial for their effective use in biotechnological applications. These insights into the crystal packing and its influence on enzyme behavior provide a valuable foundation for developing more robust and stable enzymes for industrial purposes, ensuring better performance under demanding environmental conditions. This area of structural biology is vital for understanding how molecular interactions control enzymatic activity. By identifying these interactions at the atomic level, researchers can target specific enzyme sites for regulation.

Identifying specific residues that inhibit tetramer formation in *M. metallidurans* sheds light on the molecular determinants of quaternary structure. This study aligns with prior research on structural factors influencing enzyme stability and function (<u>Sendker et al., 2024</u>). It identified specific amino acids that disrupt tetramer formation, leading to altered enzyme kinetics and stability. These findings are pivotal for protein engineering, where manipulating quaternary structure can yield enzymes with enhanced or novel properties <u>(Liu et al. 2020</u>). The *in silico* constructed tetrameric models provide a computational perspective on protein stability, complementing experimental studies. These models offer a cost-effective and rapid method for exploring structural dynamics, which is increasingly valuable in enzyme engineering <u>(Broom et al. 2020</u>, <u>Varun S. Nair et al. 2018</u>). Notably, the ECR^Mm_Dim^ may have evolved to avoid tetramerization due to environmental or functional pressures.

The ECR^Mm_Dim^ may have evolved to avoid tetramerization due to environmental or functional pressures. Collectively, these insights offer a molecular explanation for the distinct oligomeric states of *M. metallidurans* compared to other species, such as *B. ambifaria* and *K. setae*, and provide a hypothetical framework for understanding the evolutionary adaptations of these enzymes under different conditions. By elucidating how environmental pressures influence oligomerization, this study connects structural adaptations to enzymatic functionality across diverse ecological settings. Computational approaches can accelerate the development of proteins with desired characteristics for industrial or therapeutic applications, making them a vital tool in modern biotechnology (<u>Sneha Thomas et al. 2018</u>).

Comparing the primary structures and structural variations of *M. metallidurans* and *K. setae* ECRs offers valuable insights into how even subtle variations in key catalytic or stability-related residues can lead to significant functional divergence between species. This comparative evolutionary approach is essential for understanding the complex ways in which enzymes adapt to different ecological niches. The impact of these structural variations should not be underestimated; even small changes, particularly those in regions critical for enzyme activity, can result in substantial alterations in enzyme function and stability. This principle is evident in various biological systems, where minor modifications to the primary structure can profoundly influence protein behavior <u>(Stites et al. 1997</u>). Furthermore, studies have demonstrated that such variations are particularly impactful for enzymes that have evolved under different environmental conditions, as these variations can directly affect both protein stability and enzymatic activity (<u>Hartley et al. 1979</u>). In this context, understanding the influence of structural modifications on enzyme function can open up new avenues for the development of biotechnological tools and therapeutic agents based on these naturally occurring enzymes. These insights have potential applications in a wide range of scientific and industrial fields, including drug development and enzyme optimization for industrial processes (<u>Maria-Solano et al. 2018).</u>

The analysis of conformational changes in the active sites of these enzymes, particularly in response to NADPH binding, offers critical insights into the mechanisms that regulate enzymatic specificity and efficiency. The observed conformational flexibility in the active sites of ECR^Mm_Dim^ and ECR^Ks_Tet^ underscores the dynamic nature of enzyme catalysis. Conformational flexibility is often a key determinant of enzyme specificity and efficiency, as demonstrated by various studies on enzyme dynamics (<u>Dalziel et al. 1975</u>). Small structural changes can significantly impact substrate binding and reaction rates, which is essential for designing enzymes with tailored catalytic properties. This flexibility could also inform bioengineering efforts by enabling the modification of substrate specificity or reaction pathway preferences, providing valuable applications in biocatalysis (<u>Liu et al., 2020</u>).

## 4. Conclusion

The study provides valuable insights into the differences between the ECR^Mm_Dim^ and ECR^Ks_Tet^. It shows that the quaternary structure of these enzymes plays a crucial role in their stability, efficiency, and adaptability to different environmental conditions. The detailed structural analysis of the ECR^Mm_Dim^, including B-factor and ellipsoidal analyses, offers important information about the protein’s stability and dynamic properties. These findings help us understand how proteins maintain their functional integrity under varying conditions, which is essential for developing stable and robust enzymes for industrial and medical use. The study also highlights the impact of crystal packing on the enzymatic activity of ECR^Mm_Dim^, emphasizing the importance of structural context in protein function.

Identifying specific residues that inhibit tetramer formation further helps us understand the molecular determinants of quaternary structure, providing potential targets for protein engineering. Additionally, the *in silico* construction of tetrameric models provides a computational approach to exploring protein stability, complementing experimental studies and enhancing our ability to design proteins with specific properties. Comparing primary structures and structural variations between *M. metallidurans* and *K. setae* ECRs emphasizes the role of evolutionary adaptation in shaping enzyme function, with implications for biotechnology and therapeutic agents.

Finally, the analysis of conformational changes in the active sites of these enzymes, particularly in relation to NADPH binding, offers critical insights into the mechanisms that regulate enzymatic specificity and efficiency. In summary, this study advances our understanding of the structural and functional relationships in extremophilic enzymes, providing a foundation for future research and potential applications in biotechnology, medicine, and industrial processes.

### 5. MATERIALS and METHODS

### 5.1. Cell lysis, protein purification, and characterization

The process of cell lysis, protein purification, and characterization were carried out to obtain crystallography-grade ultrapure protein. First, the cells are harvested by centrifugation at 3,500 rpm for 45 minutes at 4°C. To lyse the pellet was resuspended in a lysis buffer containing 50 mM Tris-HCl pH=8, 1 M NaCl, 5% glycerol, and 100 μl Triton X-100 (Sigma-Aldrich) per 100 ml final buffer volume. The suspension underwent three rounds of sonication at 60% amplitude for 30 seconds each to further break down the cells using a Branson W250 Sonifier (USA). After sonication, the suspension was ultra-centrifuged at 35,000 rpm for 1 hour and 15 minutes at 4°C to collect the soluble fraction.

The soluble fraction was further purified using a 10 ml Ni-NTA column setup with an AKTA Go FPLC system. To ensure the column was ready, it was washed with 2 column volumes of HisA loading buffer (50 mM Tris-HCl pH=8, 300 mM NaCl, 10 mM imidazole) for equilibration. However, initial attempts at His-tag purification failed due to protein precipitation during column application. To address this, 1 M L-proline (Sigma-Aldrich) was added to the lysis and HisB elution buffers to keep the protein soluble. Once the protein was soluble, the soluble fraction was applied to the Ni-NTA column and eluted using HisB elution buffer containing 50 mM Tris-HCl pH=8, 300 mM NaCl, and 500 mM imidazole. The eluted fractions were collected using a fraction collector, and purity was analyzed via SDS-PAGE to ensure that only the target protein was present. These fractions were pooled and concentrated to 10 mg/ml using Millipore Amicon Ultra 30KDa molecular weight cut-off concentrators. Each process step was performed to ensure the final product’s high purity, necessary for subsequent applications.

To determine the ratio of dimers to tetramers in the obtained protein, size exclusion chromatography was conducted. Equal amounts of known tetrameric proteins from *M. metallidurans and B. ambifaria* were passed through an S200 column, and upon comparing the resulting peaks, it was determined that the *M. metallidurans* protein existed in a dimeric state.

### 5.2. Crystallization of *M. metallidurans* ECR

A total of 72 well sitting-drop crystallization plates (Terasaki) were prepared and subjected to screening against a library of 3500 crystallization conditions sourced from 1612 conditions from Molecular Dimensions and 1456 conditions from Hampton Research, with 432 conditions from Rigaku. In each crystallization well, a mixture of 0.83 µl from diverse crystallization buffers and 10 mg/ml of *M. metallidurans* ECR protein in a solution containing 500 mM imidazole, 300 mM NaCl, and TRIS-HCl pH 8.5 was prepared.

To slow down the crystallization kinetics, each well was hermetically sealed using 16.6 µL of 100% paraffin oil (Hampton Research). Following a one-week incubation period, crystals of the apo ECR protein manifested in diverse morphologies. Initial crystallization conditions were derived from screening conditions including Crystal Screen II, Natrix I, Wız Cryo II, PGA-LM-II, and Wizard I (Molecular Dimensions, Hampton Research). Subsequently, utilizing these solutions, plates were reconstituted, and phase separation was discerned in select wells. In instances where phase separation was observed, new plates were prepared by modulating the protein concentration under identical conditions. After that, to optimize the crystals formed in these solutions, adjustments were made to the protein quantity while maintaining a constant solution volume, i.e., plates were established with 0.83 µL of condition. Plates were inspected at biweekly intervals, culminating in crystal observation after one month. Finally, replica plates were generated utilizing the proportions of the crystals obtained.

### 5.3. Data collection, processing and structure determination

Diffraction data were collected at the Stanford Synchrotron Radiation Lightsource (SSRL) beamline 12-2 using an EIGER 2XE 16M PAD detector. The crystal belonged to the space group C121, with unit cell dimensions: a = 187.99 Å, b = 104.31 Å, c = 44.81 Å, α = 90°, β = 91.6°, and γ = 90°.

The collected data were processed using *XDS* software (<u>Kabsch, 2010</u>). The structure was solved by molecular replacement using *MOLREP* (<u>Vagin & Teplyakov, 1997</u>) with the coordinates from the apo enzyme (PDB ID: 6NA3^APO_K.setae^) as the search model. Buccaneer was used for automated model building (<u>Cowtan, 2006</u>), and manual model building was performed using *COOT* (<u>Emsley & Cowtan, 2004</u>). Refinement of the structure was carried out using the *REFMAC* program (<u>Murshudov et al., 1997</u>). Details of the data collection and refinement are presented in Table 1.

## Author Contributions

The project was initiated and coordinated by C.K. and H.D. Experimental design and data analysis were carried out by C.K., I.I.M., Y.Y., C.F., S.W., and H.D. Protein expression and purification were performed by C.K. Crystallography experiments and data collection were supported by I.I.M., S.W., and C.F. Data interpretation and manuscript writing were carried out by C.K., I.I.M., Y.Y., C.F., S.W., and H.D. All authors read and approved the manuscript. The work (proposal: 10.46936/10.25585/60000921) conducted by the U.S. Department of Energy Joint Genome Institute (https://ror.org/04xm1d337), a DOE Office of Science User Facility, is supported by the Office of Science of the U.S. Department of Energy operated under Contract No. DE-AC02-05CH11231.

## Data Availability Statement

## Supporting Information

Figures S1-7

## Acknowledgments

The authors would like to dedicate this manuscript to the memory of Dr. Albert E. Dahlberg and Dr. Nizar Türker. We sincerely thank Abdullah Kepçeoğlu and Fatma Betül Ertem for their invaluable assistance in supporting the logistical needs of this project. We are also deeply grateful to the team at Stanford University’s Stanford Synchrotron Radiation Lightsource (SSRL). Use of the Stanford Synchrotron Radiation Lightsource, SLAC National Accelerator Laboratory, is supported by the U.S. Department of Energy, Office of Science, Office of Basic Energy Sciences under Contract No. DE-AC02-76SF00515. The SSRL Structural Molecular Biology Program is supported by the DOE Office of Biological and Environmental Research, and by the National Institutes of Health, National Institute of General Medical Sciences (P30GM133894). The contents of this publication are solely the responsibility of the authors and do not necessarily represent the official views of NIGMS or NIH.

## Notes

### Competing Interest Statement

The authors have declared no competing interest.

