## Supplementary material for "A Novel Unorthodox Dimeric Primary Enoyl-CoA Reductase Structure": supplemantary figure1

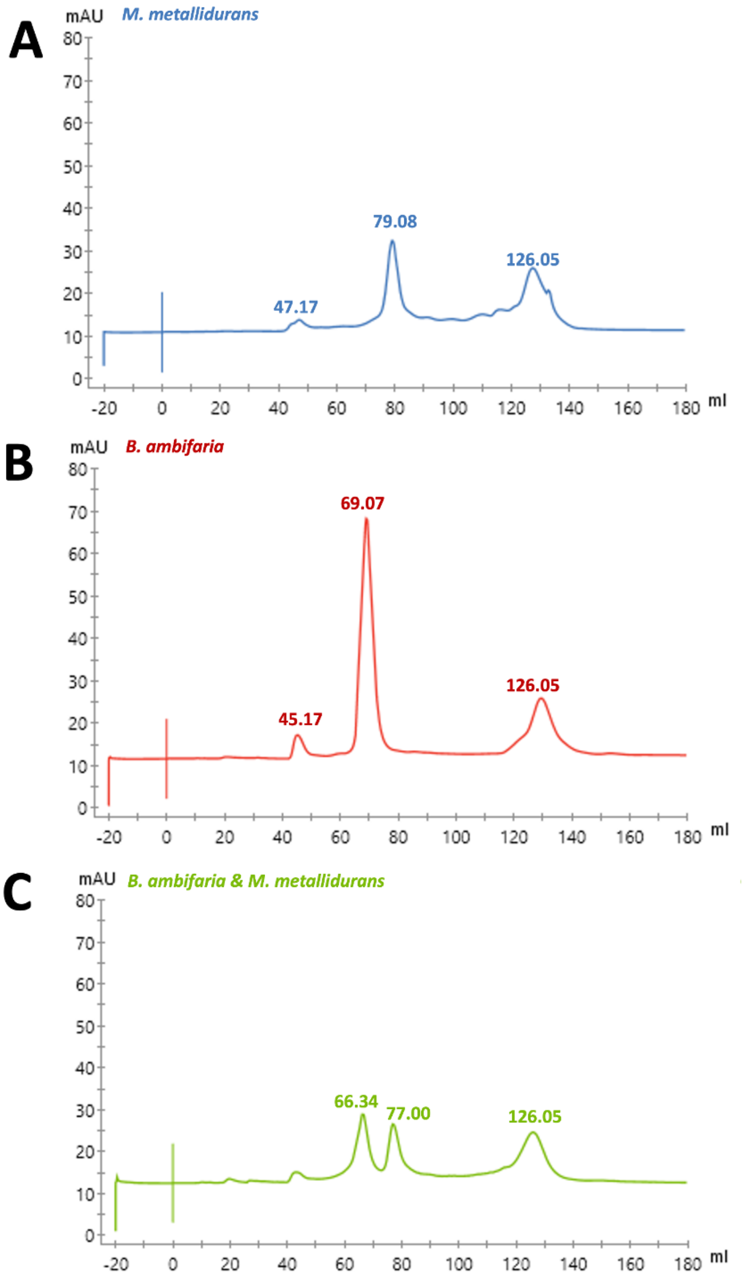


**Supplementary Figure 1: Size exclusion chromatography profiles**, the following size exclusion chromatography profiles illustrate the behavior of *B. ambifaria* and *M. metallidurans* proteins, both individually and when mixed: **a)** Size exclusion chromatogram of a 1 mL pure sample of *M. metallidurans* protein (blue). The dimer peak was observed at 79.08 mL. **b)** Size exclusion chromatogram of a 1 mL pure B.ambifaria protein (red) sample. The tetramer peak was observed at 69.07 mL. **c)** Size exclusion chromatogram of a mixture containing 500 µL of *B.ambifaria* protein and 500 µL of *M. metallidurans* protein (total volume 1 mL) (green). The tetramer peak of *B. ambifaria* protein was observed at 66.34 mL, the dimer peak of *M. metallidurans* protein was observed at 77.00 mL, and an additional elution volume is observed at 126.05 mL.
