## Supplementary material for "A Novel Unorthodox Dimeric Primary Enoyl-CoA Reductase Structure": supplemantary figure3

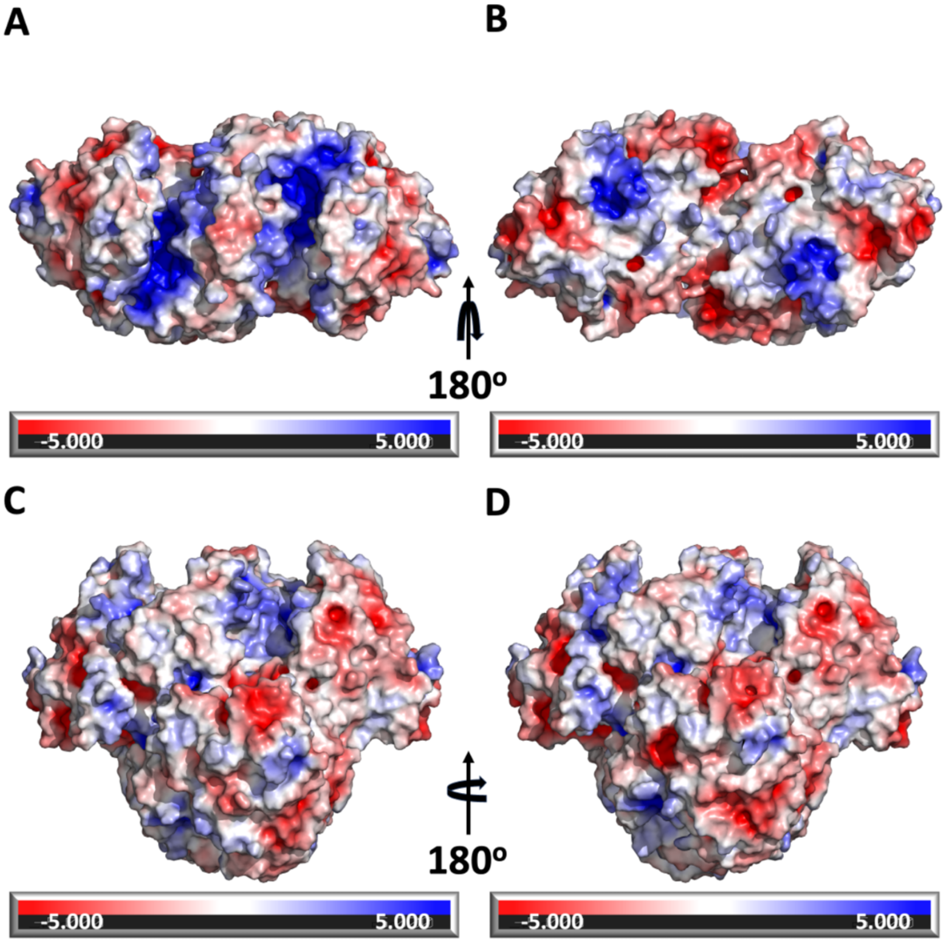


**Supplementary Figure 3: Electrostatic surface potential comparison of *M. metallidurans* and *K. setae* ECR (Enoyl-CoA carboxylase/reductase) structures.** **a)** and **b)** show the electrostatic surface potential of the *M. metallidurans* dimer from two opposite orientations, rotated by 180°. The distribution of positive (blue) and negative (red) electrostatic regions highlights key surface features of the dimer. **c)** and **d)** represent the electrostatic surface potential of the *K. setae* tetramer, also rotated by 180° to display both views. The tetramer reveals distinct electrostatic characteristics compared to the dimer, emphasizing the differences in surface charge distribution between the dimeric and tetrameric forms. The color bar indicates the electrostatic potential scale, ranging from -5.000 (red) to +5.000 (blue) kT/e.
