## Supplementary material for "A Novel Unorthodox Dimeric Primary Enoyl-CoA Reductase Structure": supplemantary figure4

**A**

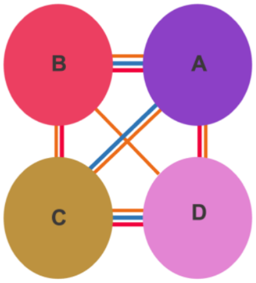


AlphaFold predictions of *M. metallidurans* tetramer

| Chains | No. of interface residues | Interface area $(A^2)$ | No. of salt bridges | No. of disulphide bonds | No. of hydrogens bonds | No. of non-bonded contacts |
|---|---|---|---|---|---|---|
| A=B | 36:36 | 1800:1801 | 4 | - | 24 | 302 |
| C=D | 36:36 | 1807:1802 | 4 | - | 24 | 310 |
| A=C | 2:2 | 152:156 | - | - | 1 | 3 |
| A=D | 2:2 | 113:115 | 2 | - | - | 16 |
| B=C | 2:2 | 141:139 | 2 | - | - | 24 |
| B=D | 1:1 | 101:98 | - | - | - | 1 |

**B**

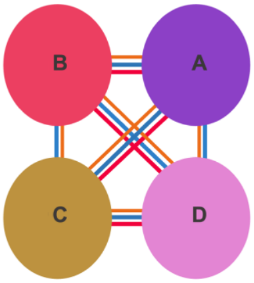


*K. setae* PDB ID:6NA3 APO

| Chains | No. of interface residues | Interface area $(A^2)$ | No. of salt bridges | No. of disulphide bonds | No. of hydrogens bonds | No. of non-bonded contacts |
|---|---|---|---|---|---|---|
| A=B | 20:20 | 955:954 | 4 | - | 16 | 86 |
| C=D | 20:20 | 942:942 | 4 | - | 16 | 86 |
| A=C | 35:34 | 1599:1605 | 2 | - | 22 | 259 |
| B=D | 35:35 | 1596:1590 | 2 | - | 21 | 256 |
| A=D | 19:18 | 1126:1122 | - | - | 12 | 118 |
| B=C | 19:19 | 1124:1128 | - | - | 12 | 119 |

**Supplementary Figure 4: Comparison of the interface characteristics between the AlphaFold-predicted tetrameric structure of *M. metallidurans* ECR and the experimental structure of *K. setae* ECR tetramer. a)** Interface analysis of the *M. metallidurans* ECR tetramer from AlphaFold prediction. The schematic on the left shows the arrangement of the four chains (A, B, C, and D) in the tetramer with colored lines representing different types of chain interactions. The table on the right provides detailed interface metrics for each chain pair, including the number of interface residues, interface area (Å²), salt bridges, disulfide bonds, hydrogen bonds, and non-bonded contacts. **b)** Interface analysis of the experimental structure of *K. setae* ECR tetramer (PDB ID: 6NA3, Apo form). The schematic and table layout are similar to panel (A), showing the interactions between the chains based on the experimental structure. The comparison highlights differences in interface areas and the number of interactions between the AlphaFold prediction and the experimental data, particularly in hydrogen bonds and non-bonded contacts.
