## Supplementary material for "A Novel Unorthodox Dimeric Primary Enoyl-CoA Reductase Structure": supplemantary figure5

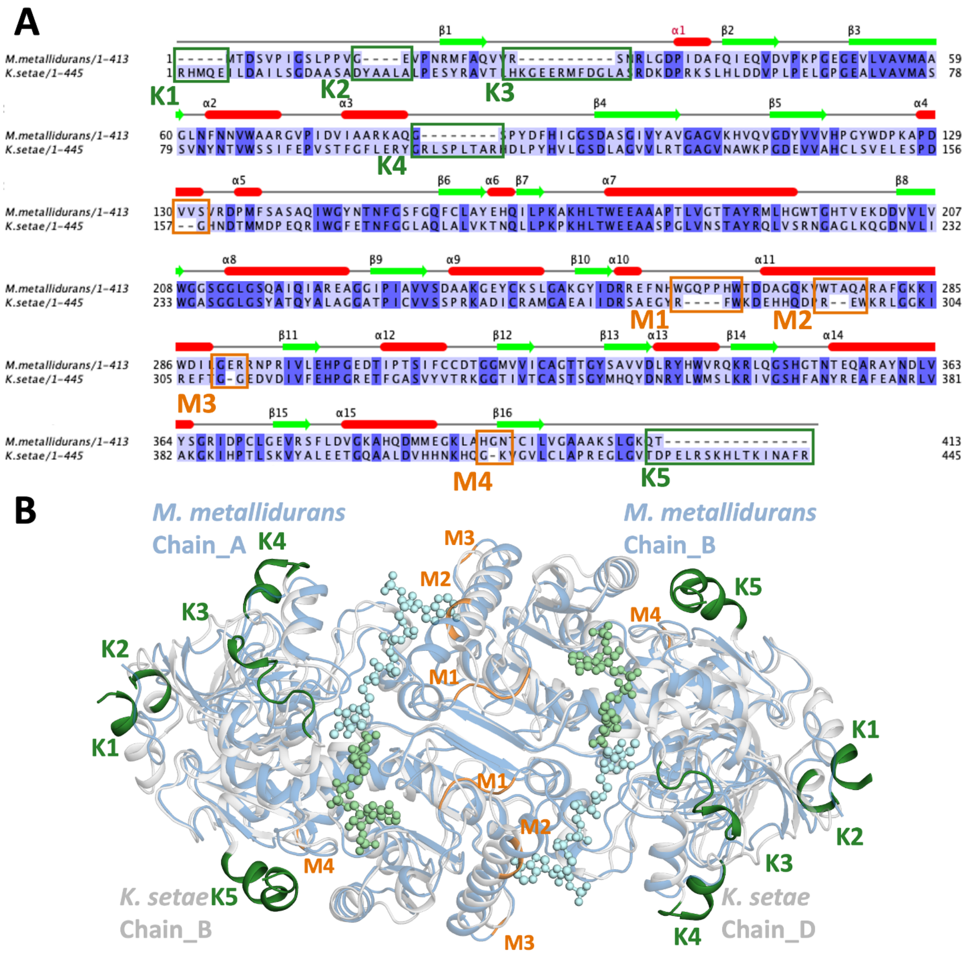


**Supplementary Figure 5: Comparative sequence and structural analysis of ECRs from *M. metallidurans* and *K. setae*,** **a)** Sequence alignment of ECRs from *M. metallidurans* (1–413) and *K. setae* (1–445). The secondary structural elements of the *M. metallidurans* ECR are annotated above the sequences, with helices represented as red cylinders and beta strands as green arrows. Key regions that differ between the two sequences are highlighted. Differences in the *M. metallidurans* structure compared to *K. setae* are labeled as K1-K5 (green), while differences in *K. setae* compared to *M. metallidurans* are labeled as M1-M4 (orange). **b)** Structural superposition of the *M. metallidurans* dimer (cyan) and the open subunits of the apo *K. setae* tetramer (blue-white). Regions that differ in the *M. metallidurans* structure compared to *K. setae* are highlighted and labeled as K1-K5 (green), while regions that differ in *K. setae* compared to *M. metallidurans* are labeled as M1-M4 (orange). These variations are shown to impact the overall structural differences between the two ECRs.
