## Supplementary material for "A Novel Unorthodox Dimeric Primary Enoyl-CoA Reductase Structure": supplemantary figure6

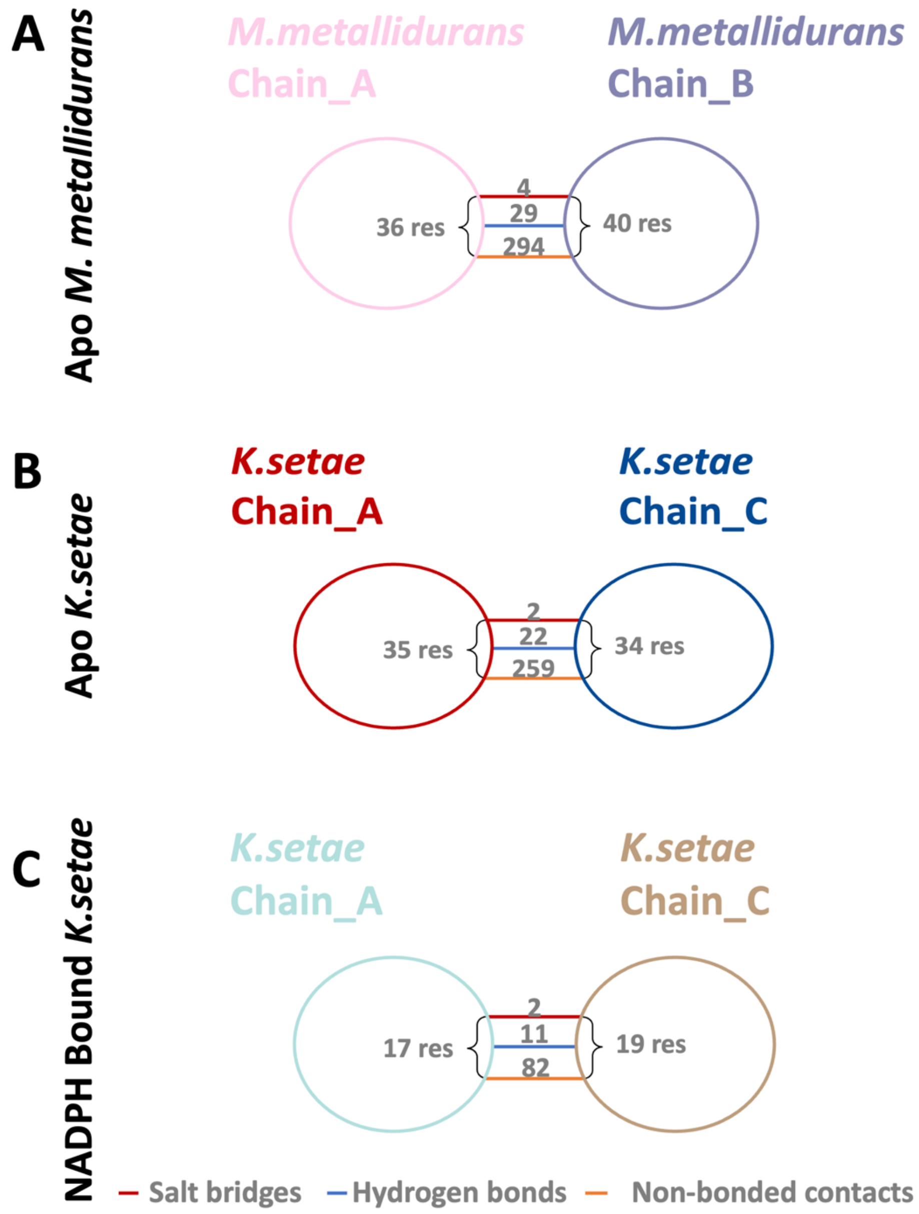


**Supplementary Figure 6: Structural Analysis of Dimer and Tetramer Interfaces**, **a)** Shows the numbers of salt bridges, hydrogen bonds, and non-bonded contacts between homologs in the Apo *M. metallidurans* dimer structure. You can see these interactions in the interaction residues given in Scheme 1. **b)** Shows the number of salt bridges, hydrogen bonds, and non-bonded contacts in the dimer portion of the Apo *K. Setae* tetramer structure (PDB ID: 6NA3). You can see these interactions in the interaction residues given in Scheme 2. **c)** Shows the number of salt bridges, hydrogen bonds, and non-bonded contacts in the dimer portion of the NADPH Bound *K. setae* tetramer structure (PDB ID: 6NA6). You can see these interactions in the interaction residues given in Scheme 3.

**Note:** Each panel details the interactions between two chains (chain_A and chain_B or chain_C) and categorizes them into salt bridges (red), hydrogen bonds (blue), and non-bonded contacts (orange). Additionally, the number of residues involved in the contact surface for each chain is indicated.
