## Supplementary material for "A Novel Unorthodox Dimeric Primary Enoyl-CoA Reductase Structure": supplemantary figure7

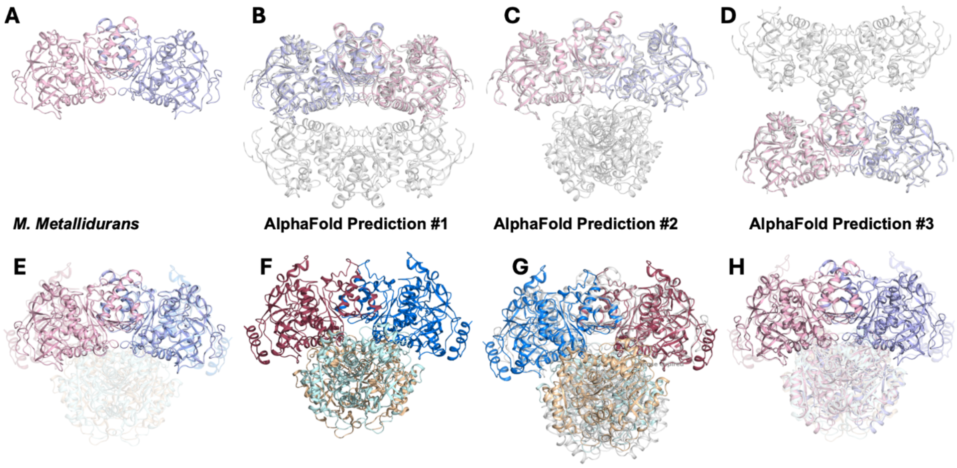


**Supplementary Figure 7: Comparison of the *M. metallidurans* ECR dimer with AlphaFold tetramer predictions and alignment with *K. setae*. a)** Crystal structure of the ECR dimer from *M. metallidurans*, with each chain colored pink and blue. **b)** AlphaFold Prediction #1 for the ECR dimer, showing a symmetrical arrangement with two dimers stacked on each other. **c)** AlphaFold Prediction#2 displays a more compact tetramer configuration, with the dimers rotated relative to each other. **d)** AlphaFold Prediction #3 reveals an extended arrangement in which the dimers interact through different interface regions compared to Predictions #1 and #2. **e)** Structural alignment of the *M. metallidurans* ECR dimer and the *K. setae* apo form. **f)** Tetrameric representation of the apo form of *K. setae*. **g)** Superposition between AlphaFold Prediction #2 and the apo form of the *K. setae* tetramer. **h)** Superposition of the ECR dimer structure alongside the in-silico modeled tetramer of *M. metallidurans* ECR.
